# Efficient genome editing in multiple salmonid cell lines using ribonucleoprotein complexes

**DOI:** 10.1101/2020.04.03.022038

**Authors:** Remi L. Gratacap, Ye Hwa Jin, Marina Mantsopoulou, Ross D. Houston

## Abstract

Infectious and parasitic diseases have major negative economic and animal welfare impacts on aquaculture of salmonid species. Improved knowledge of the functional basis of host response and genetic resistance to these diseases is key to developing preventative and treatment options. Cell lines provide a valuable model to study infectious diseases in salmonids, and genome editing using CRISPR provides an exciting avenue to evaluate the function of specific genes in those systems. While CRISPR/Cas9 has been successfully performed in a Chinook salmon cell line (CHSE-214), there are no reports to date of editing of cell lines derived from the most commercially relevant salmonid species Atlantic salmon and rainbow trout, which are difficult to transduce and therefore edit using lentivirus-mediated methods. In the current study, a method of genome editing of salmonid cell lines using ribonucleoprotein (RNP) complexes was optimised and tested in the most commonly-used salmonid fish cell lines; Atlantic salmon (SHK-1 and ASK cell lines), rainbow trout (RTG-2) and Chinook salmon (CHSE-214). Electroporation of RNP based on either Cas9 or Cas12a was efficient at targeted editing of all the tested lines (typically > 90 % cells edited), and the choice of enzyme expands the number of potential target sites for editing within the genomes of these species. These optimised protocols will facilitate functional genetic studies in salmonid cell lines, which are widely used as model systems for infectious diseases in aquaculture.

## Introduction

Salmonid fish are amongst the highest value aquaculture species globally, together worth in excess of $22Bn in 2017 (FAO 2019). However, infectious disease outbreaks are a continuous threat to sustainable production and future expansion to meet global demands for these fish. Therefore, development of vaccines and therapeutics is an important goal, and selective breeding for improved host resistance has major potential to help tackle several diseases (Yáñez et al. 2014). Genomic selection has also been applied to enhance the rate of genetic gain for disease resistance traits in breeding programmes (Houston 2017; Zenger et al. 2019), and genome editing approaches may offer further step-improvements in the future (Gratacap et al., 2019b). However, research into the functional mechanisms underlying host response to salmonid pathogens, and host genetic variation in resistance is important to support development of these potential solutions.

Genome editing using CRISPR systems is a valuable research tool because it allows targeted changes to genomes of species of interest. Therefore, CRISPR editing can be applied to test the functional role of a particular gene or variant in a trait of interest, such as resistance to infection (Staller et al. 2019). This can be achieved by editing the target species’ genome at a location that will result in knockout of the target gene, or by introducing domains which will activate or repress its expression (Gilbert et al. 2014; Doudna and Charpentier 2014). CRISPR/Cas9 has successfully been applied *in vivo* to edit the genome of Atlantic salmon (Edvardsen et al. 2014) and rainbow trout (Cleveland et al. 2018), including to create a sterile salmon by knockout of the *dnd* gene (Wargelius et al. 2016). In addition to understanding gene function, genome editing holds significant potential to be applied in commercial aquaculture to tackle major production barriers (Gratacap et al., 2019b).

The use of cell lines for research into salmonid pathogens and host response has been well-established, and engineering of those cell lines holds substantial promise for advancing fish health research (Collet et al. 2018). However, genome editing of salmonid cell lines remains in its infancy, with the first report of successful CRISPR editing being in the Chinook salmon (*Oncorhynchus tshawytscha*) cell line (CHSE-EC, derived from CHSE-214), which was engineered to stably express Cas9 and EGFP (Dehler et al. 2016). This line has subsequently been applied to develop a clonal STAT2 knockout line to study the role of this gene in viral response (Dehler et al. 2019), and as a proof-of-principle to demonstrate that transduction of lentivirus facilitates high-efficiency editing (Gratacap et al. 2019a). Additionally, Escobar-Aguirre et al. (2019) reported delivery and expression of a gRNA, Cas9, and an mCherry reporter gene in CHSE-214 using a plasmid construct. However, other salmonid fish cell lines are considered difficult to transfect and to develop clonal lines (Collet et al. 2018), making analogous approaches in existing Atlantic salmon and rainbow trout cell lines challenging. Indeed lentivirus transduction using the approach of Gratacap et al. (2019a) resulted in a very low success rate in the Atlantic salmon SKH-1 cell line (<1 % of cells successfully transduced; data not shown). CRISPR/Cas9 ribonucleoprotein (RNP) complexes have potential for editing of fish cell lines, as demonstrated with efficiency of up to 62 % in medaka (*Oryzias latipes*) (Liu et al. 2018). Furthermore, Cas12a editing has been successfully applied in mammalian cells, *Xenopus* and zebrafish, including using RNP systems (Moreno-Mateos et al. 2017; Liu et al. 2019), and significantly expands the number of ‘editable’ sites in the target species’ genomes. The efficiency of Cas9 or Cas12a editing using RNP in salmonid cell lines is unknown, and these approaches may help overcome the aforementioned challenges to cell line editing in cell lines of Atlantic salmon and rainbow trout; two of the world’s most important aquaculture species.

In the current study, a simple and reproducible method of editing multiple salmonid fish cell lines using electroporation of Cas9 RNP complexes is presented. The method was tested and optimised, resulting in very efficient editing of all the commonly used salmonid cell lines tested; specifically, Atlantic salmon (ASK and SHK-1), rainbow trout (RTG-2) and Chinook salmon (CHSE-214). Additionally, electroporation of Cas12a RNP which uses a different protospacer adjacent motif (PAM) of 5’TTTV led to high genome editing (although less than Cas9), expanding the number of potential target editing sites in these species’ genomes.

## Materials and Methods

### Cell lines

The cell lines used in this study were: (i) salmon head kidney 1 (SHK-1), an immortalised cell line from Atlantic salmon (*Salmo salar*) obtained from the European Collection of Authenticated Cell Cultures (ECACC) (97111106); (ii) Atlantic salmon kidney (ASK), an immortalised cell line from Atlantic salmon (*S. salar*) obtained from American Type Culture Collection (ATCC; CRL-2747); (iii) rainbow trout gonad (RTG-2), an immortalised cell line from rainbow trout (*O. mykiss*) obtained from ECACC (90102529), and (iv) Chinook salmon embryo 214 (CHSE-214), an immortalised cell line from Chinook salmon (*O. tshawytscha*) obtained from ECACC (91041114). All cells were grown as a monolayer in L15 media (Sigma-Aldrich, St. Louis, USA) supplemented with heat inactivated fetal bovine serum (FBS) (Gibco, Waltham, USA) (SHK-1, 5 %; RTG-2, CHSE-214 and ASK, 10 %), 40 μM β-mercaptoethanol (Gibco) for SHK-1, 100 Units/mL penicillin and 100 μg/mL streptomycin (Gibco). All cells were cultured in an incubator at 22 ± 1 °C without CO_2_. SHK-1 was split 1:2 at 80 % confluency with ⅓ conditioned media and the rest of cell lines were split 1:3 – 1:4 with fresh media.

SHK-1 cells were used for the initial optimisation of RNP editing, as described below. As part of this process, an SHK-1 line with a GFP transgene was created to allow testing of gRNA targeting knockout of this transgene causing loss of fluorescence. This cell line (SHK-fuGFP) was generated by transfecting a CMV-GFP_Puromycin construct (Addgene 45561, a gift from Michael McVoy) in SHK-1 cells with Fugene HD transfection reagent (Promega, Madison, USA). To achieve this, SHK-1 cells were plated in a 24-well plate at 40,000 cells per wells and incubated overnight at 22 °C. Media was replaced with 500 μL of L15 (10 % FBS, no antibiotics) containing 0.5 μg plasmid and 1.5 μL of FugeneHD (ratio Fugene:DNA 3:1, according to manufacturer’s instructions). After 7 days, cells were selected with Puromycin at a concentration of 1 μg/mL for a period of 4 weeks.

### Optimisation of RNP transfection and editing

To test and optimise the RNP platform, an intergenic region of the Atlantic salmon genome (Genbank accession NC_027325.1 ssa26; 15004350-15004900) was targeted with a gRNA via transfection of the SHK-1 cell line. This was followed by validation of the optimised conditions by EGFP knockout in the SHK-FuGFP cell line and knockout of coding region of *slc45a2* (Gene ID: 106563596).

The crRNAs were designed with CRISPOR (http://crispor.tefor.net/) and the CRISPR Design Tool (Synthego Inc, Menlo Park, USA), and crRNAs and tracrRNAs were ordered from IDT (details of all gRNA are given in Table 1). The RNP complexes were assembled as follows: crRNA and tracrRNA were resuspended in nuclease free water at 100 μM, aliquoted and frozen at −80 °C. One μL of crRNA and 1 μL of tracrRNA were mixed and incubated at 95 °C for 5 min. The mixture was cooled to room temperature and 2 μL of 20 μM Cas9 (NEB, Ipswich, USA) was added (final concentration of 10 μM of Cas9 and 25 μM of gRNA). The complexes were incubated at room temperature for 15 min and kept on ice until use.

**Table 1.**
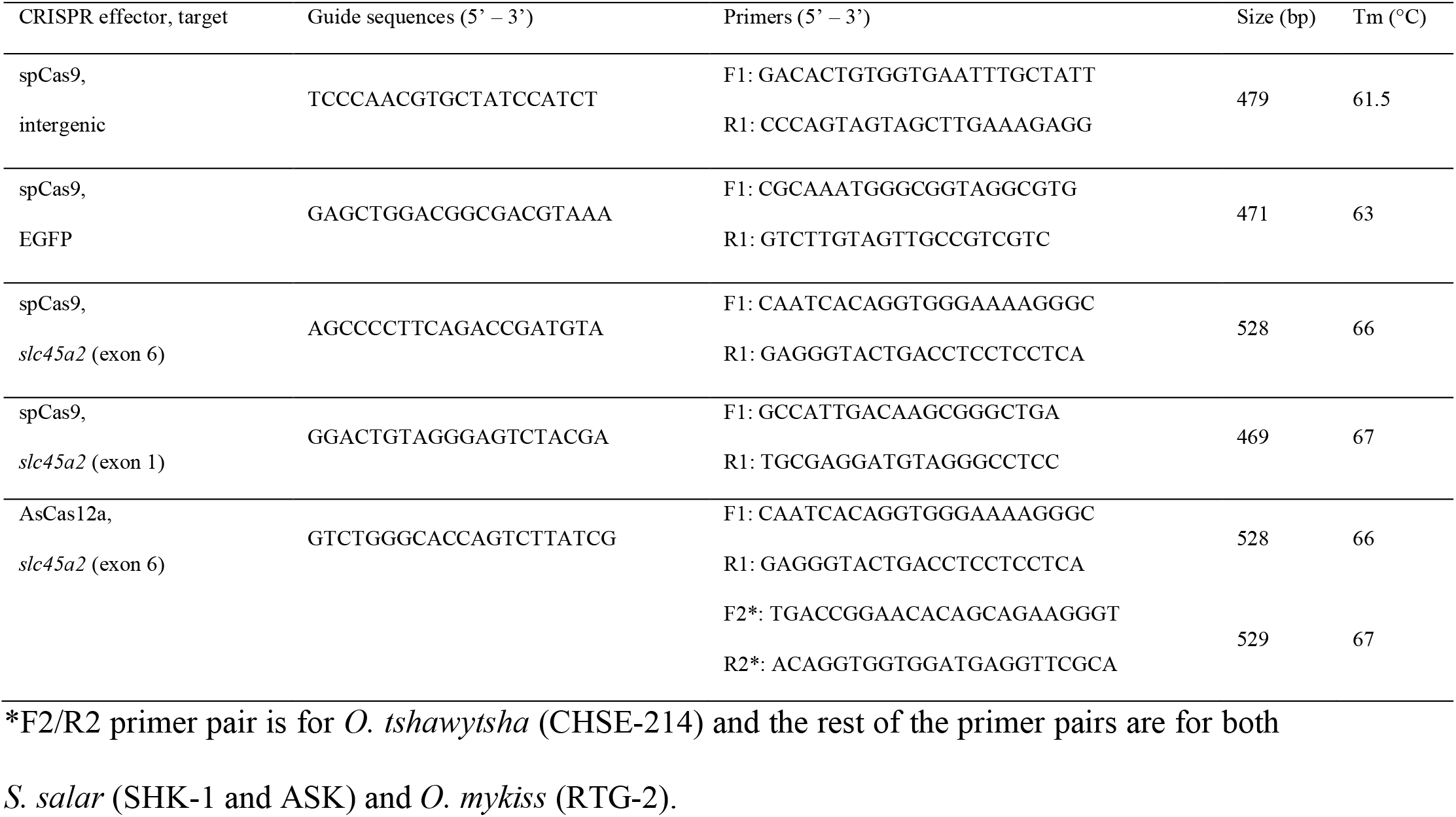
Guide RNA sequences and primers used for amplification and sequencing of target genomic regions

The first optimisation step involved varying the concentration of the Cas9-gRNA RNP complex, with the starting point being electroporation conditions that have previously been successful in plasmid transfection of SHK-1 (data not shown). To achieve this, different concentrations of RNP were diluted in OptiMEM reduced serum media (Gibco) (final volume 4 μL) and mixed with 10 μL of SHK-1 cells at 107 cells/mL in OptiMEM (final concentrations ranges from 0.0875 to 2.8 μM Cas9 RNP). After 5 min incubation at room temperature, the cells plus the RNP were electroporated with the Neon system (Invitrogen, Carlsbad, USA) according to the manufacturer’s instructions but with OptiMEM instead of Neon R Buffer (Invitrogen). Briefly, cells were centrifuged at 600g for 4 min and resuspended in 1 mL of OptiMEM. Cells were counted using a hemocytometer and adjusted to 10_7_ cells/mL by centrifugation/resuspension in the appropriate volume of OptiMEM. An aliquot of 4 μL of the RNP complex was added to 10 μL of cells suspension (105 cells total) which was then incubated for 5 min at room temperature. The mixture was then electroporated using 10 μL tips and dispensed in 1 mL of fresh media in a 24-well plate. 100 μL of the suspension was transferred to a 96-well plate (104 cells) for genomic DNA isolation or cell culture. The cells were incubated overnight at room temperature, and the media changed to 1 mL of fresh media. Once the cells reached confluency (in 24-well plates), they were resuspended in media (using trypsin) and divided 1:2 (adding 33 % conditioned media for the SHK-1 cells). Cells were kept in 96- or 24-well plates for gDNA isolation at 1, 2, 4, 7 and 14 days post treatment (dpt) or expanded to 6-well plates once they reached confluency for measurement of fluorescence using flow cytometry at 14 dpt.

### Cell survival

In addition to assessing the transfection and editing efficiency of the SHK-1 cells, the cell viability was tested in parallel for each of the setting used for the optimisation protocol using CellTiter-Glo 2.0 (Invitrogen). In brief, following electroporation, 100 μL of the cells in the 24-well plate were transferred to a 96-well plate and incubated for 48 hours. Surviving cells still attached to the bottom of the plate were rinsed once in PBS and 120 μL of CellTiter-Glo solution (diluted 1:10 in PBS) was added to each well. The plate was incubated in the dark for 30 min on a plate rocker at room temperature and 100 μL of the solution was transferred to a flat bottom white wall 96-well plate (Greiner Bio-One, Austria). The luminescence was measured using a Cytation3 imaging reader and the Gen5 software V3.03 (BioTek, Winooski, USA).

### Validation of optimised RNP editing by GFP knockout

A second test of RNP editing was performed in the SHK-fuGFP cells by using the optimised settings to transfect an RNP complex with a gRNA targeting knockout of the GFP transgene. Following the transfection, the loss of GFP was measured by flow cytometry. To achieve this, the cells were trypsinized and resuspended in PBS. The cells were kept on ice and flow cytometry was performed using a Fortessa-X20 (BD Biosciences, San Jose, USA). Single-cell events were gated and the percentage of GFP-positive cells and the intensity of GFP fluorescence from each cell was measured.

### Assessing the efficiency and nature of the edits using Sanger sequencing

Testing of editing efficiency was performed by isolation of genomic DNA followed by PCR amplicon Sanger sequencing. Genomic DNA (gDNA) was extracted with QuickExtract buffer (Lucigen, Middleton, USA) by adding 30 μL to a well of a 96-well plate and incubating for 5 min. The samples were then processed according to the manufacturer’s instructions (65 °C for 15 min and 98 °C for 2 min). PCR was performed with 50 μL reactions using NEB Q5 and 1 μL of the gDNA with 33 cycles amplification at optimal annealing temperature (Table1). 5 μL of the PCR product was run on a 1.5% agarose gel to verify correct amplification. Amplified sequence was purified with AmPURE XP magnetic beads (Agencourt, Beverly, USA) according to the manufacturer’s instructions (using 1:1 ratio) and sent to GATC/Eurofins (Germany) for Sanger sequencing. Analysis of the chromatograms (based on.abi files) was used to assess the editing efficiency and nature of the induced edits using the Inference of CRISPR Edits (ICE, Synthego Inc) software for Cas9 or TIDE software (Brinkman et al. 2014) for Cas12a to determine the editing efficiency (% of cells containing putative indels).

### Testing of RNP editing in other salmon cell lines

Following optimising of the electroporation and incubation settings for the SHK-1 cells described above, similar protocols were tested in the other three cell lines by targeting the *slc45a2* gene. Several combinations of different electroporation settings (1200 – 1600 V, 10 – 40 ms, 1 – 3 pulses) and cell resuspension buffers (Neon R buffer and OptiMEM) were tested to achieve highest transfection rate using tracrRNA-ATTO550 (IDT, Coralville, USA) by detecting ATTO550-positive cell population using flow cytometry at 24 hours post electroporation. The best transfection result (99.9 – 100 %) was obtained with 1400 V 20 ms 1 pulse for RTG-2 and 1600 V 10 ms 3 pulses for CHSE-214 and ASK with OptiMEM as a resuspension buffer (data not shown).

The Cas9 RNP complex was assembled as described above and Cas12a RNP was formed by adding 31.2 pmol of AsCas12a (IDT) and 50 pmol of crRNA (IDT) per 10_5_ cells. The complexes were incubated at room temperature for 15 min and kept on ice until use. The final concentration of 1 μM of Cas9 RNP and 2.6 μM of AsCas12a RNP were tested with the optimised electroporation settings for each cell line. At 7 dpt the editing efficiency and the nature of the induced edits were assessed by Sanger sequencing and ICE software for Cas9 or TIDE software for Cas12a as described above.

## Results

### Electroporation of Cas9-gRNA complex leads to efficient editing

To test and optimise the CRISPR Cas9 genome editing platform using RNP, an intergenic region of the Atlantic salmon genome was targeted in the SHK-1 cell line. The first optimisation step involved varying the concentration of the Cas9 RNP complex, with set electroporation conditions (previously optimised for plasmid transfection of SHK-1: 1300 V, 30 ms and 1 pulse). The editing efficiency increased with increasing concentration of RNP up to 1.4 μM, but plateaued at higher concentrations (**Fig.1a**). This RNP concentration was then used for optimisation of electroporation settings. Both cell survival and editing efficiency (**Fig.S1a** and **Fig.S1b**) were assayed. Using three pulses of 1600 V for 10 ms resulted in the highest editing rate of 42 % at 4 days post treatment (dpt, **Fig.1b**), and also the highest cell survival (113 % survival compared to control, **Fig.S1b**). To assess whether editing was still occurring after four days, samples of the SHK-1 cells were taken for Sanger sequencing at 7 and 14 dpt. The proportion of edited cells was higher after 7 days than 4 days, but did not increase afterwards (**Fig.1b** and **Fig.S1c**). The resulting optimised RNP editing protocol for SHK-1 cells, led to 57 % of the cells edited using electroporation of 1.4 μM RNP with three 1600 V pulses of 10 ms, followed by incubation for 7 days. It is worth noting that the edit pattern (+1, −1 and −6 bp edits) generated by this gRNA:Cas9 complex is very reproducible as seen in Fig S1D.

**Figure 1:**
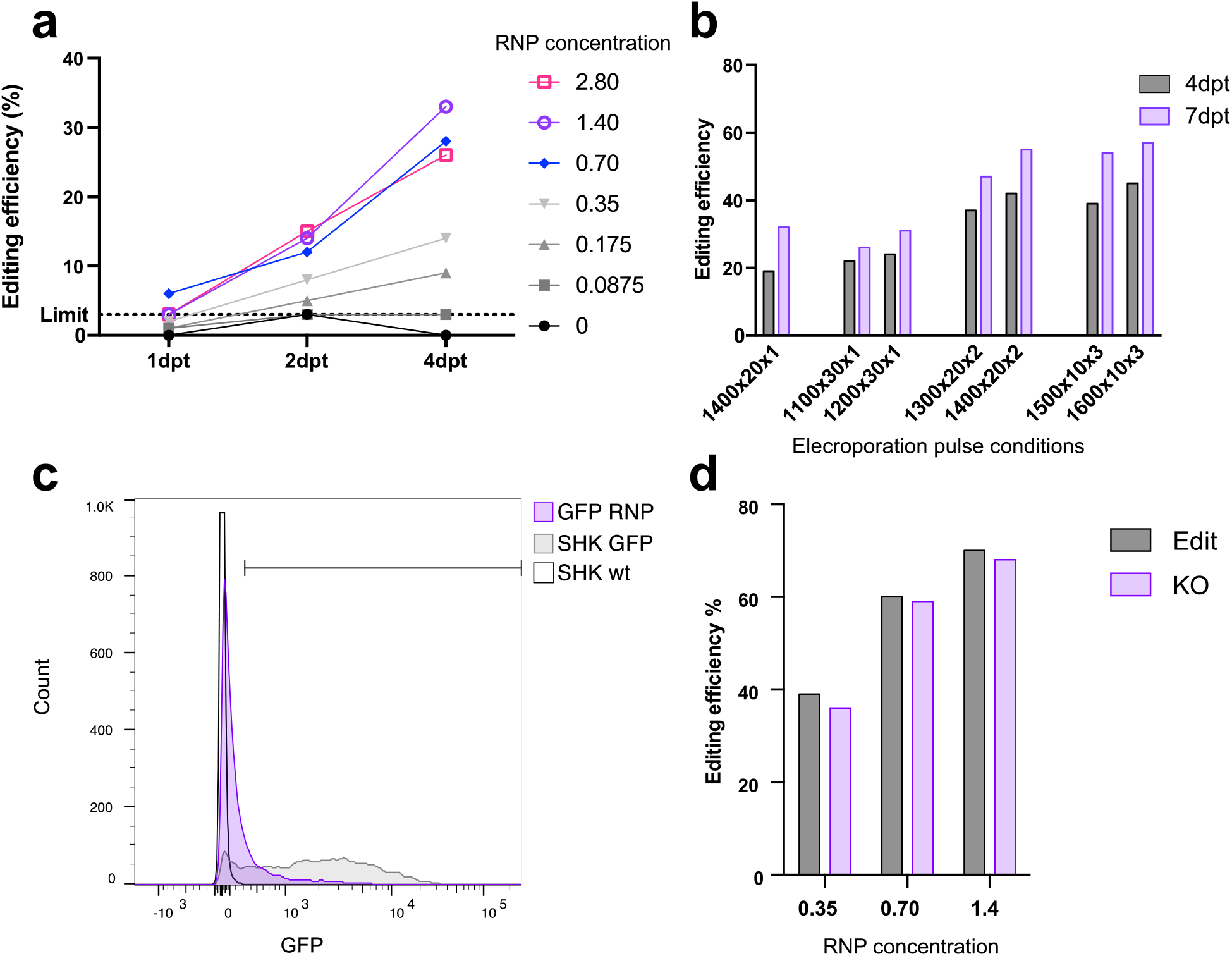
Efficient editing of Atlantic salmon cell line by electroporation of Cas9 RNP. **Fig.1a and 1b**. Optimisation of genome editing in SHK-1 cells targeting an intergenic region. SHK-1 cells were electroporated (1300 V, 30 ms and 1 pulse) with different concentrations of Cas9 RNP (μM) and gDNA isolated at different timepoints after electroporation (days post transfection, dpt). **Fig.1b**. Using the optimal concentration (1.4 μM) of RNP, different electroporation settings were evaluated [voltage (V) x pulse duration (ms) x number of pulses] and the sampling time increased to 4 and 7 dpt. **Fig.1c and 1d**. Efficient knock out of GFP in SHK-GFP cells. SHK-fuGFP were electroporated with optimised settings and gRNA targeting GFP transgene. After 14 days, fluorescence was measured using flow cytometry (**Fig.1c**), and editing efficiency was also assessed by Sanger sequencing (**Fig.1d**) as well as predicted transcript knockout (purple bars). All genome editing efficiency generated using ICE (Synthego Inc) deconvolution of Sanger sequencing chromatogram.

Further validation of the Cas9 RNP electroporation platform was performed using an EGFP knockout system in SHK-1 cells. An SHK-1 cell line with constitutive EGFP expression was created, and an RNP complex targeting EGFP was designed. Using the optimised electroporation and RNP concentration established above, there was approximately 75 % loss of GFP as measured by flow cytometry (**Fig.1c**), with an estimated 68 % editing efficiency by Sanger sequencing (**Fig.1d**).

### Optimised protocol translates to efficient editing in multiple cell lines using Cas9 and Cas12a enzymes

To evaluate the potential of RNP editing in the most commonly used salmonid cell lines, the optimised settings established were used to target a coding region of *slc45a2* which is involved in pigmentation and has been successfully knocked out using CRISPR in Atlantic salmon (Edvardsen et al. 2014). The cell lines targeted were SHK-1 and ASK (*S. salar*), RTG-2 (*O. mykiss*) and CHSE-214 (*O. tshawytsha*). Electroporation of Cas9 RNP targeting *slc45a2* resulted in over 90 % of cells edited in SHK-1, RTG-2 and ASK and over 70 % edited in CHSE-214 (**Fig.2a** and **Fig.2b**), which was consistent across two independent experiments. It is worth noting that the same gRNA was used in all cell lines but there was a 1 bp mismatch with the homologous target region of Chinook salmon (CHSE-214), which may explain the lower editing observed.

**Figure 2:**
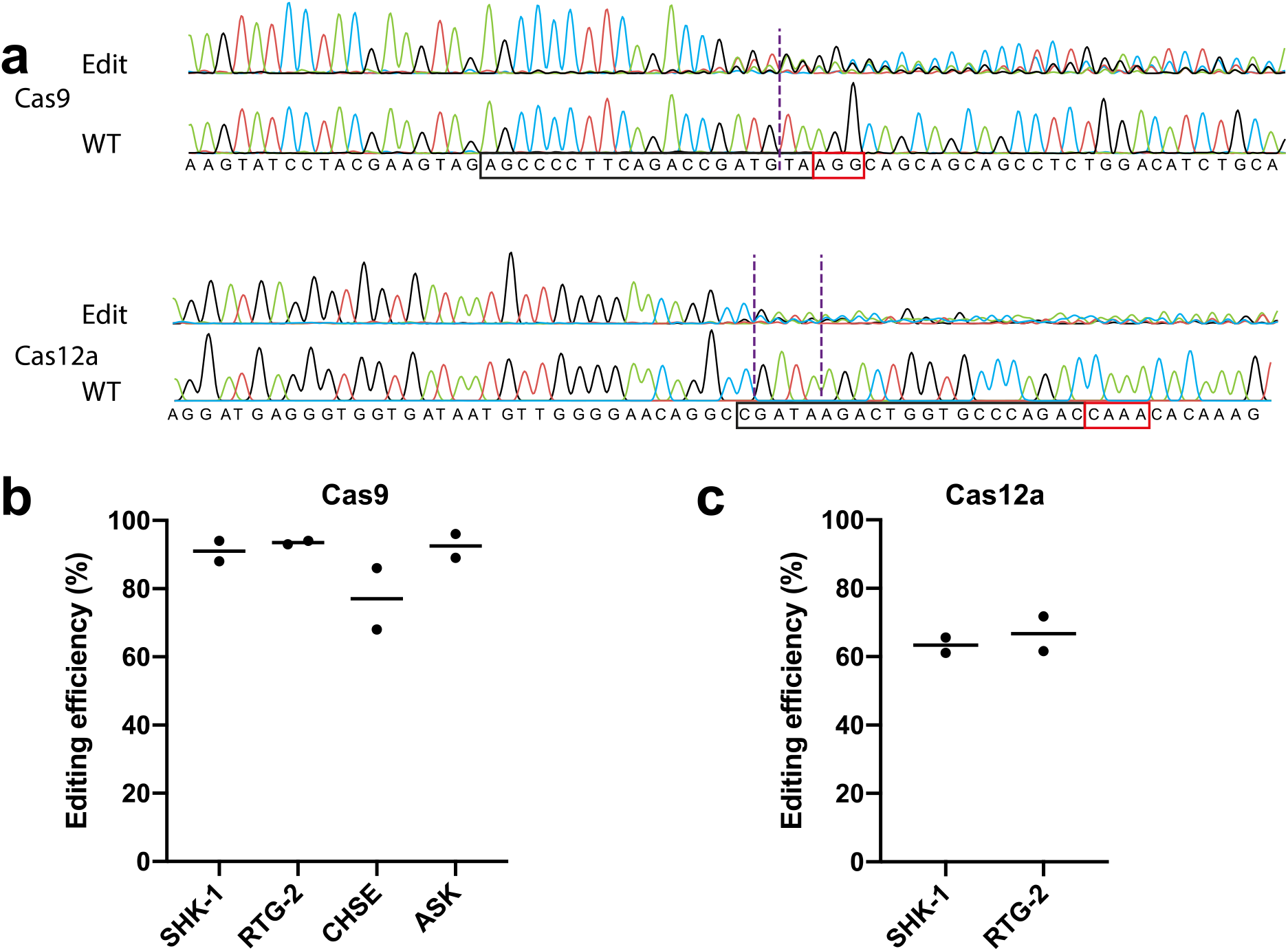
Efficient editing of salmonid cell lines with different Cas proteins. **Fig.2a**. Representative chromatogram of the Sanger sequencing for the target region of the *slc45a2* gene in SHK-1 cells, either wild-type (WT) or edited with Cas9 (top) and Cas12a (bottom) RNP. The binding regions are boxed in black and PAM sequence is in red. The nuclease cut positions are indicated by dashed lines. **Fig.2b**. Editing of *slc45a2* gene in SHK-1, ASK, RTG-2 and CHSE-214 using Cas9 RNP. **Fig.2c**. Editing of *slc45a2* gene in SHK-1 and RTG-2 with Cas12a RNP. Two independent experiments (with median) are represented. Editing efficiency was estimated with ICE and TIDE analyses for Cas9 and Cas12a RNP, respectively.

The nature of the edits made to the genome of in the target region was assessed using Sanger sequencing of PCR amplicons followed by analysis using the ICE software (as described above). In both experiment replicates, the majority of SHK-1 cells edited by Cas9 contained a 1 bp deletion, whereas the same gRNA primarily resulted in a 1 bp insertion in RTG-2 cells (**Fig.S2b**). While the pattern of indels observed when using the same gRNA varied between cell lines, single base pair insertions or deletions were predominant (**Fig.S2b**).

The Cas12a RNP targeted the same genomic region as for Cas9 RNP (*slc45a2* Exon 6 Ssa01:117877060-117877079 for Cas9 and Ssa01:117877013-117877033 bp for Cas12a), and Cas12a also resulted in high editing efficiency in both Atlantic salmon (SHK-1, 63 %) and rainbow trout (RTG-2, 67 %) cell lines (**Fig.2c**), although the editing rates were notably lower than when using the Cas9 enzyme.

## Discussion

The current study presents an optimised method for genome editing in the most commonly used salmonid cell lines using electroporation of Cas9 or Cas12a RNP. The method can be used to edit over 90% of cells in a mixed cell line population with Cas9, and over 60% with Cas12a; this is a higher editing efficiency than reported in medaka fish cell lines using Cas9 RNP (62 % editing, Liu et al., 2018). This method circumvents several challenges to genetic engineering of many fish cell lines due to their slow growth, poor transfection/transduction efficiency, and difficulty to obtain clonal lines (Collet et al., 2018).

In the current study, electroporation was optimised by testing a wide range of voltage (850 – 1600 V), time (10 – 40 ms) and number of pulse (1 – 3) variables, and the most effective electroporation setting for SHK-1 was shown to be with 1600 V 10 ms 3 pulses. Interestingly, these settings were also optimal for CHSE-214 and ASK, while the optimal electroporation setting for RTG-2 was 1400 V 20 ms 1 pulse. This is the first report of electroporation of RNP in salmonid cells and might explain the difference in optimal electroporation settings compared with previous studies using plasmids (Ojima et al. 1999; Chi et al. 2012; Marivin et al. 2015). These settings were also optimal for electroporation and editing using Cas12a RNP in both Atlantic salmon and rainbow trout cell lines. The use of Cas12a as well as Cas9 increases the range of targetable sequences for editing (5’NGG and 5’TTTV) in salmonid species’ genomes.

Interestingly, editing efficiency in the SHK-1 cell line was higher at 7 dpt than 4 dpt. This implies that editing is occurring more slowly than in mammalian systems (Kim et al. 2014) and highlights that the Cas9 protein is still active for over a week in the experimental conditions described (cells incubated at room temperature). It is also noteworthy that both gRNAs targeting *slc45a2* were more efficient than the gRNA targeting the intergenic region. This highlights that the efficacy of the system will vary across different target genomic regions, which may be due to differences in chromatin accessibility. Therefore, certain genes and genomic locations may not be amenable to highly efficient editing using this approach. This problem also applies to lentivirus and plasmid delivery systems, but low efficiency editing in these systems can be combined with selection using antibiotics or fluorescence to enrich for edited cells, which is not possible using the RNP editing system described herein. Ultimately, single-cell cloning might be required to achieve a high level of editing (possibly 100 % edited clones) for sites with lower editing efficiency using the current system. However, single cell cloning has not been successful in SHK-1 cells, and the process is likely to take several months in the other salmonid cell lines due to their slow-growing nature.

By utilising Cas RNP complexes, the method presented here allows rapid and efficient editing of salmonid cell lines. From design to experimental testing of the edits, the protocol takes just over two weeks. This is approximately half the time required for lentivirus or plasmid delivery as the constructs have to be generated and sequenced before delivery and then enrichment applied. Additionally, lentivirus and plasmid approaches require investigation and optimisation of the promoter (Ruiz et al. 2008; Martinez-Lopez et al. 2013) and selection marker (Schiøtz et al. 2011) choices for in each cell line since little information is yet available for most fish cell lines. Finally, the very high editing rate (> 90 %) obtained with the method described herein also circumvents the need for enrichment of the edited population, allows direct testing of the cells for the phenotype of interest.

The proposed Cas RNP method has potential for versatile CRISPR/Cas editing applications in salmonid fish cell lines, which are widely used model systems to understand genetics and immunology of commercially and environmentally important fish species. The ability to perform targeted gene knockout will allow for assessment of candidate genes involved in genetic resistance to disease, for example. The *in vitro* system could also act as a testbed for gRNAs efficiency prior to their use *in vivo*. Given the scientific and commercial interest in salmonid fish species, and that all the cells line tested could be efficiently edited, this technique is likely to form a useful component of the toolbox for functional genetics and immunology research in fish.

---

**Supplementary Figure S1.**
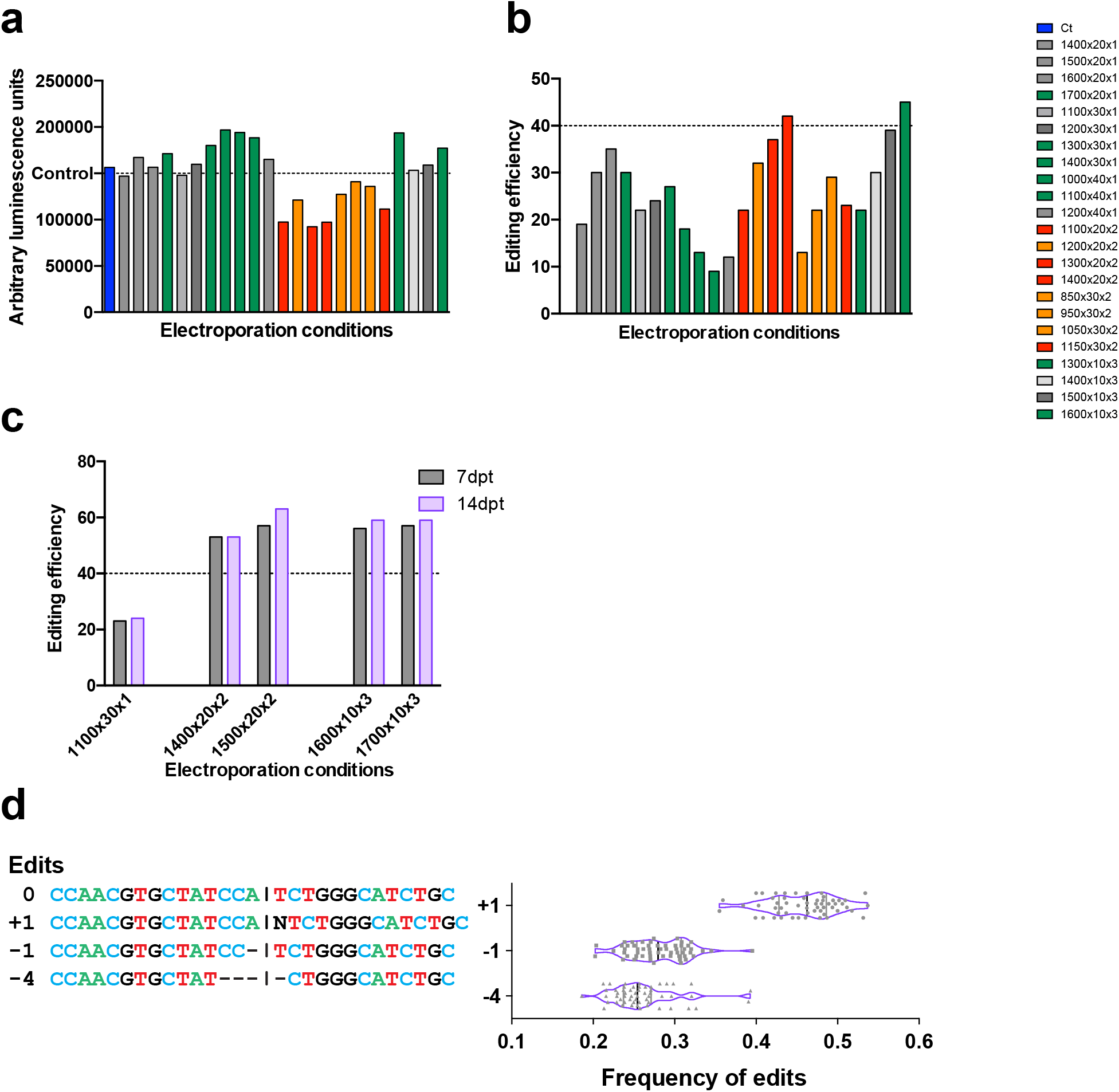
SHK-1 cells were electroporated with 1.4 μM Cas9:gRNA RNP and transferred to 2 separate 96-well plates. (A) After 48h, cell survival (plate1) was calculated using CellTiter Glo 2.0. (B) genomic DNA (plate 2) was extracted at 7 dpt, and the target sequence amplified by PCR and editing efficiency estimated using Sanger sequencing. (C) Using 1.4 μM RNP, the editing efficiency after 7 and 14 ddpt was estimated for different electroporation settings. (D) All the sequencing data, obtained from ICE analysis of Sanger sequencing of the intergenic target region from optimisation experiments (n=55) were pooled and plotted according to edit pattern.

**Supplementary Figure S2.**
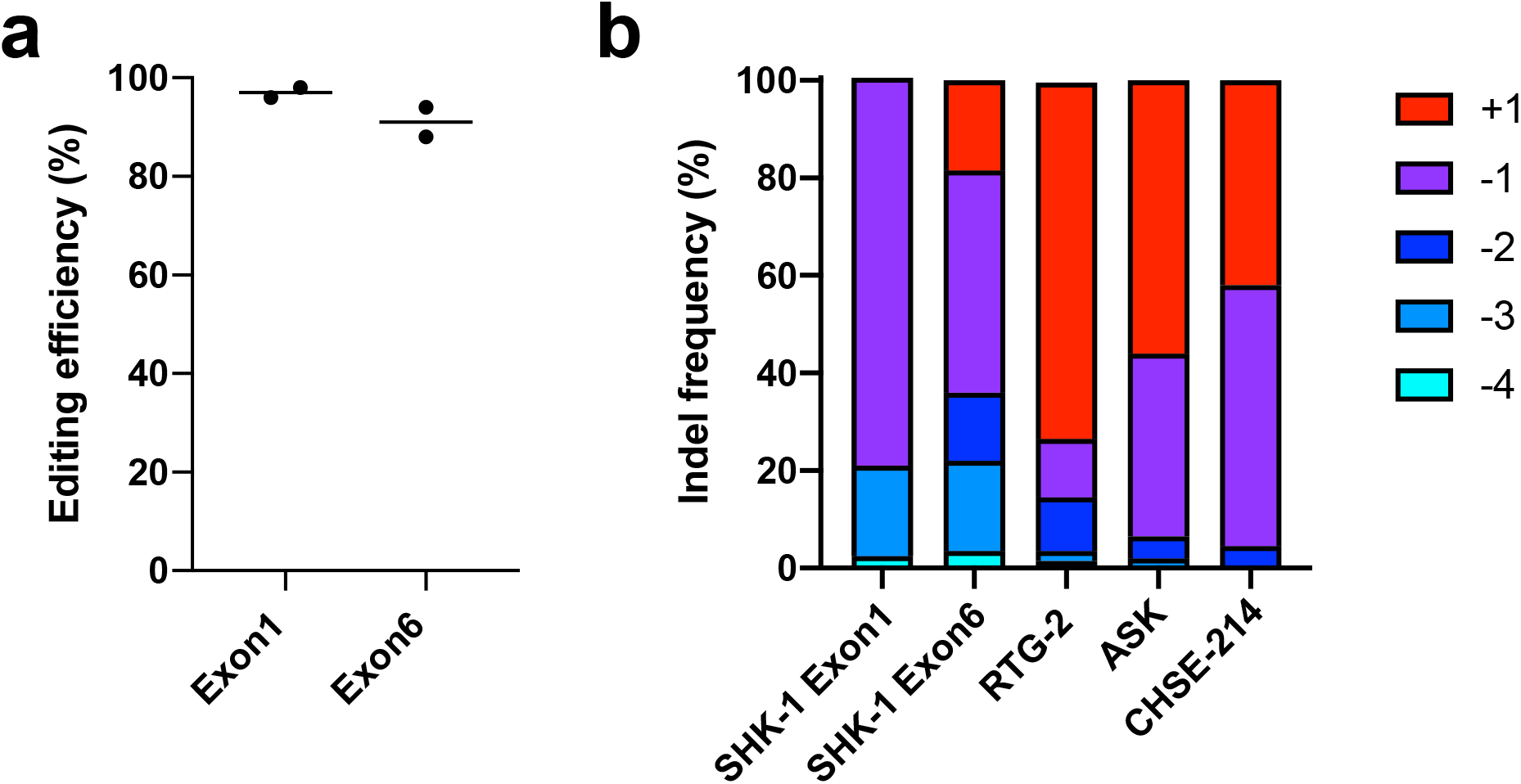
(A) Editing of *slc45a2* gene in SHK-1 with Cas9 RNP with two gRNA targeting different exons (Exon 1 and Exon 6). (B) Detail of the Indel frequency estimated for *slc45a2* with Cas9 in different cell lines (refers to Fig 2B). Indels with frequencies over 2 % are represented. Two independent experiments (with median) are represented. Editing efficiency was estimated using ICE analysis.

## Acknowledgments

The authors acknowledge funding from the Biotechnology and Biological Sciences Research Council (BBSRC), via the ‘AquaLeap’ project (BB/S004343/1) and Institute Strategic Programme grants (BBS/E/D/20002172, BBS/E/D/30002275 and BBS/E/D/10002070). The authors also acknowledge support and funding from Hendrix Genetics.

## Declaration

### Availability of data and material

The raw data that support the findings of this study and the material described are available from the corresponding author on request.

### Authors contribution

Ross Houston, Remi Gratacap and Ye Hwa Jin contributed to the study conception and design. Material preparation, data collection and analysis were performed by Remi Gratacap, Ye Hwa Jin and Marina Mansopoulou. The first draft of the manuscript was written by Ross Houston, Remi Gratacap and Ye Hwa Jin and all authors commented on previous versions of the manuscript. All authors read and approved the final manuscript.

